# Considerations for performance metrics of metagenomic next generation sequencing analyses

**DOI:** 10.1101/2020.12.17.423212

**Authors:** Jason G. Kralj, Stephanie L. Servetas, Samuel P. Forry, Scott A. Jackson

## Abstract

Evaluating the performance of metagenomics analyses has proven a challenge, due in part to limited ground-truth standards, broad application space, and numerous evaluation methods and metrics. Application of traditional clinical performance metrics (i.e. sensitivity, specificity, etc.) using taxonomic classifiers do not fit the “one-bug-one-test” paradigm. Ultimately, users need methods that evaluate fitness-for-purpose and identify their analyses’ strengths and weaknesses. Within a defined cohort, reporting performance metrics by taxon, rather than by sample, will clarify this evaluation. An estimated limit of detection, positive and negative control samples, and true positive and negative true results are necessary criteria for all investigated taxa. Use of summary metrics should be restricted to comparing results of similar cohorts and data, and should employ harmonic means and continuous products for each performance metric rather than arithmetic mean. Such consideration will ensure meaningful comparisons and evaluation of fitness-for-purpose.

## Introduction

Metagenomic next generation sequencing (mNGS) workflows employ any of a variety of taxonomic classification tools and are capable of comparing DNA against databases of 1,000s or even 10,000s of organisms. As a result, clinical metagenomics have seen tremendous development in the past decade and are poised to revolutionize infectious disease diagnostics. This has advantages over traditional culture-based or PCR-based methods, which are typically highly specific but difficult to parallelize. At this point, the question is not if metagenomics analyses work, but what data and metrics are appropriate to evaluate and validate performance. However, guidance around this has been unclear.^1^ As a result, widespread acceptance of mNGS diagnostic methods has yet to meet its potential.

The performance metrics (PMs) sensitivity, specificity, precision, accuracy, etc. are the benchmark for evaluating analytical performance and fitness for purpose and have generally agreed upon definitions. Carefully setting the positive and negative controls and results are critical if PMs are to have validity and utility, especially in the clinical mNGS diagnostic space. At its heart, mNGS is a technology that enables massive parallelization of DNA-based single-organism testing; the reports/analyses summarize all the tests performed. mNGS taxonomic classifiers often perform well identifying genus- and species-level taxonomies, though the lack of consensus metrics and formats for data reporting make direct comparisons between results from different classifiers difficult.^2,3^ Further, follow-on analysis to filter results based on limit-of-detection, minimum number of mapped reads, genome coverage, coverage depth, and relative abundance cutoffs are necessary.^4^ The aforementioned filters have demonstrated improved performance of these analyses,^1^ and this study presupposes the completion of that substantial work before moving to performance analysis.

One common approach to calculate performance metrics is to sum results across all possible taxa within a database, which results in misleading specificity and accuracy scores always near 100 %. Consider that many databases have tens of thousands of species, but only a small percentage are of diagnostic interest. By summing the results across all possible taxa, an mNGS assay could entirely miss 100 of the most prevalent species and still be 99 % accurate. While some will argue the issue is with the database, we propose it is due to the approach. One way to prevent these misleading results would be to evaluate analysis performance with respect to each taxon.

Herein, we propose reporting performance metrics by taxon, which we call an *organism-centric* evaluation. Calculating and organizing PMs this way will ensure that relevant organisms can be thoroughly investigated, poor results are not overlooked, and non-diagnostic or untested taxa are excluded. Additionally, where two or more workflows are evaluated using the same raw sequencing output, a summary of the PMs across all taxa could be accomplished using the harmonic mean (HM) and continuous product (Pi, combined probabilities) for each performance metric. Unlike arithmetic mean, HM and Pi are well-suited to rate and fractional data. We propose this methodology in the context of mNGS analysis development and to aid evaluation of analysis performance and clinical utility.

## Performance metrics (PMs)

### A Sample-centric vs. Organism-centric View

It is important to consider the end user when evaluating these technologies; any positive (or negative) result for *each organism* listed in the report will be taken at face value. Hence, an analysis that has a weakness in performance for one or a few organisms impacts the credibility of that entire analysis. Any review or validation should identify such deficiencies, as they ultimately impact utility.

There is general agreement on the definitions for PMs:

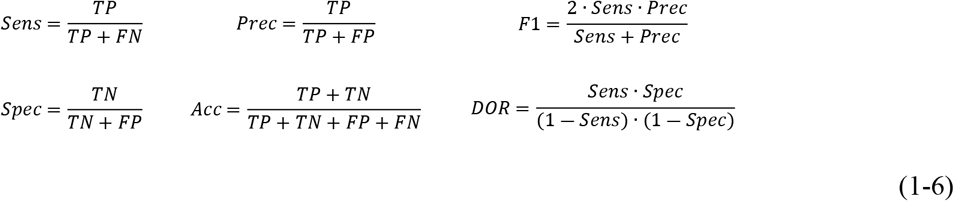

Where *Sens* is sensitivity, *Spec* is specificity, *Prec* is precision, *Acc* is accuracy, *F*1is the harmonic mean of sensitivity and precision, and *DOR* is the diagnostic odds ratio.

However, the definition of what constitutes a true/false positive/negative (TP/TN/FP/FN) within mNGS has created confusion and ambiguity for the field. mNGS assays are employed using large databases of organisms (often more than 10^4^); therefore, one common way to evaluated mNGS assays is by examining a summary analysis of performance for a sample. Thus, the current practices can be viewed as *sample-centric*, where the tallies for TP/TN/FP/FN are made across all organisms within the analysis of a single sample. Combining all organisms within a single TP or FN for each sample can lead to overlooking deficiencies at the organism level. Furthermore, this type of analysis is fundamentally flawed because it treats the mNGS analysis as a single test. On the contrary, mNGS analysis is a platform to perform highly multiplexed testing for each organism in its database. This distinction is critical because it establishes the framework for evaluation.

We propose the use of an *organism-centric* approach for evaluation. For the PMs to enable accurate evaluation an mNGS workflow, every taxon being evaluated must have TP and TN results several criteria must be fulfilled prior to starting the mNGS analysis. This requires positive (known presence) and negative controls (known absence) within samples for evaluating the performance of any assay. Importantly, for each taxon with a predefined cohort, TP and TN results are required. This presupposes (1) the inclusion of positive (known presence) and negative (known absence) controls within the sample set and (2) analytical validity for each taxon including a limit of detection (LOD) estimate. If positive or negative controls are excluded from an analysis characterization, no conclusions can be drawn about the performance of an assay for that organism.

Employing the organism-centric approach, for each taxon *i*, we can calculate the performance metric of interest (for example, sensitivity):

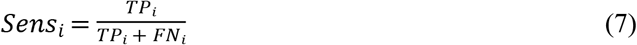

### Summary Performance Metrics

While the use of summary values can simplify evaluation, they can also lead to significant loss of information. Maintaining the context (dataset, cohort) of evaluation is critical. Using different datasets with the same mNGS analysis can produce different PMs; thus, making comparison of *different* analyses is invalid if the cohort is not controlled.^2^

When a comparison is appropriate, the harmonic mean (HM) of PMs, instead of the arithmetic mean (AM), provides more useful information because PMs are rates. This highlights potential deficiencies such as outliers. When combining across all organisms/taxa tested we can estimate the average performance for a taxon. For *n* organisms:

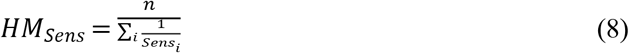

We propose including the product of each PM across all organisms (Pi) to help identify how deficiencies compound to impact the report, considering any one thing misidentified or omitted lowers the confidence in the entire analysis.

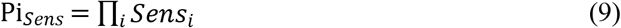

Either small numbers of large deficiencies or large numbers of small deficiencies would manifest in a low score. Pi values approaching 1 indicate few or no deficiencies in an assay with respect to that performance metric. The stringency of the Pi metric may not be necessary to validate many applications, but would serve as a powerful differentiator between analyses.

Thus, we propose the reporting of results with the following reporting format:

1. A brief tabulated list of lowest-performing organisms including TP/FP/TN/FN and performance metric values. (The full table should be appended to any report in an easily-readable format, such as comma-separated or tab-delimited)
2. A summary score for each performance metric using harmonic mean (HM) and product (Pi), if warranted.

## Example Datasets and Analyses

### An *Organism-centric* Report

Consider a hypothetical example where we are studying 10 different sample mixtures, with each sample containing 5/10 possible organisms. We calculated the individual performance metrics, as well as HM and Pi values. The exact identities, taxonomic classifier, and postprocessing analysis used do not matter here.

Using an *organism-centric* report, ordered to highlight low performers, we can quickly identify organisms creating challenges and decide on a course of action (**Table 1,** *Analysis Results*). Faced with this, developers/users would likely consider their options, such as update or upgrade the database or change an LOD/threshold in an earlier data processing step. If these do not improve the PMs, this test may be unreliable in assessing organisms *Ak, Bl*, and *Cm*. Summary performance metrics (HM & Pi) provide a means to evaluate this analysis against others especially if considering an extensive list of organisms; with a small cohort they are unnecessary.

**Table 1.** Proposed summary analysis of a hypothetical dataset using 10 samples with combinations of 10 organisms. In this example, organisms Ak, Bl, & Cm show relatively poor performance, and may need to be further studied depending on the application.

| Analysis Results |  |  |  |  | Performance Metrics |  |  |  |  |  |  |
| --- | --- | --- | --- | --- | --- | --- | --- | --- | --- | --- | --- |
| Organism | TP | FP | TN | FN | n | Sens | Pr | Spec | Acc | F1 | DOR |
| *Ak* | **3** | **4** | **1** | **2** | 10 | **0.6** | **0.43** | **0.2** | **0.4** | **0.5** | **0.4** |
| *Bl* | **2** | 1 | 4 | **3** | 10 | **0.4** | **0.67** | 0.8 | **0.6** | **0.5** | 2.7 |
| *Cm* | **3** | 1 | 4 | **2** | 10 | **0.6** | **0.75** | 0.8 | **0.7** | **0.67** | 6.0 |
| *Dn* | 4 | 1 | 4 | 1 | 10 | 0.8 | 0.8 | 0.8 | 0.8 | 0.8 | 16.0 |
| *Eo* | 5 | 1 | 4 | 0 | 10 | 1 | 0.83 | 0.8 | 0.9 | 0.91 | 26.7 |
| *Fp* | 5 | 0 | 5 | 0 | 10 | 1 | 1 | 1 | 1 | 1 | 100 |
| *Gq* | 5 | 0 | 5 | 0 | 10 | 1 | 1 | 1 | 1 | 1 | 100 |
| *Hr* | 5 | 0 | 5 | 0 | 10 | 1 | 1 | 1 | 1 | 1 | 100 |
| *Is* | 5 | 0 | 5 | 0 | 10 | 1 | 1 | 1 | 1 | 1 | 100 |
| *Jt* | 5 | 0 | 5 | 0 | 10 | 1 | 1 | 1 | 1 | 1 | 100 |
| Summary Metrics |  |  |  |  |  | Sen | Pr | Spec | Acc | F1 | DOR |
| Harmonic Mean |  |  |  |  |  | **0.76** | **0.79** | **0.67** | **0.77** | **0.78** | 6.5 |
| Product |  |  |  |  |  | **0.12** | **0.14** | **0.08** | **0.12** | **0.13** | **0.01** |

Summary Metrics give a limited insight into average and total performance, but may be necessary when evaluating large cohorts and/or different analysis tools. The harmonic mean estimates average individual taxon performance, and the product serves to compound a metric across the cohort, effectively scoring the overall report. Because these Summary Metrics strongly depend on context (cohort size and analysis setpoints), using them to predict performance with other cohorts or applications should be done with caution.

The table was organized with the lowest performing organisms first. Values below 80 % or significantly different from others were made BOLD to demonstrate how a performance threshold could be applied. DOR values were limited to 100.

In contrast, if the *sample-centric* approach is applied to the same 10 samples, it can result in misleading PMs. If we conservatively estimate 1000 taxa in the database, we would report *Sens*=0.8, *Prec*=0.8, *Spec*=0.992, and *Acc*=0.984. While you may reevaluate your analysis given these results, unlike the *organism-centric* approach, it is hard to identify the reasons for these scores. Expanding the database to 10^4^ by including irrelevant organisms artificially improves the results for *Spec* and *Acc* to 0.9992 and 0.9984, respectively, further masking the relatively poor performance with respect to *Ak, Bl*, and *Cm*.

The *organism-centric* approach could be modified to suit a variety of purposes, including real-world samples where hundreds or thousands of organisms are being considered, or where tuning around subspecies or strain-level identification is required. The generation of performance metrics even for large cohorts requires little computational power. Before any physical experiments are performed, *in silico*-generated mock datasets could rapidly examine an analysis pipeline and identify appropriate experimental conditions, numbers of samples, and/or breadth of cohort needed.

### Reevaluation of a Well-Reasoned Study

In another example, we evaluated data from previous work,^5^ where they report on the performance of an analysis of cell-free DNA for 1250 organisms. We chose to highlight this study because the authors included proper controls and provided comprehensive testing of both *in silico* and clinical samples with sufficient information to enable evaluation of the results. What we propose would improve the transparency of reporting and highlight potential deficiencies that may be deleterious to overall confidence in an analysis.

In the Specificity Analysis, the authors reported specificity per analyte of 99.998 %, a precision of 99.2 % to 99.4 % per sample from simulated samples, a 92.1 % precision from simulated clinical samples, and sensitivity of 93.6 % from simulated clinical samples. Each value was generated using the arithmetic mean across all samples and organisms within a cohort. The 3 cohorts were (a) an asymptomatic patient plasma pool (all negative controls), (b) a panel of 1250 simulated high-level spike-in controls containing one of each organism, and (c) 125 simulations of cell-free DNA near the LOD. With the data provided,^5^ we generated results containing the proposed performance metrics for each cohort (**Table 2**).

**Table 2.**
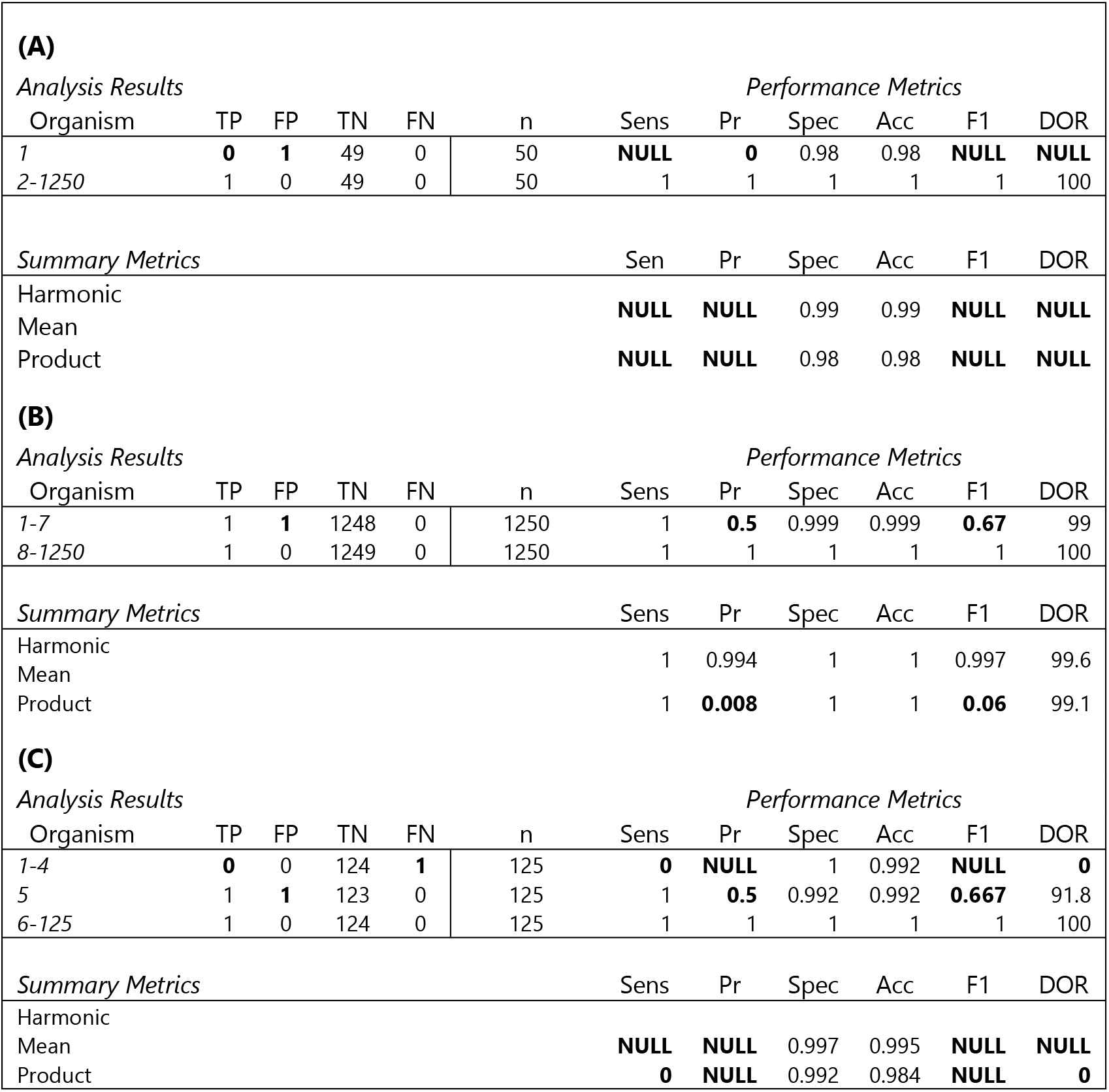
Three cohorts from the Blauwkamp (2019) study. (A) Results from the negative control experiment (healthy plasma) have no TP, and so performance metrics cannot be determined. (B) The results from high-abundance in silico spike-in experiments show TP and TN for every taxon examined. 7 samples have a false positive, leading to precision scores of 50 % for those organisms. Using additional positive control samples may show that the precision is higher, or could indicate that these organisms have high incidence of FP. (C) Simulated in silico experiments show results from near-LOD clinical indicate 4 without a TP, meaning sensitivity and precision cannot be determined. It may be possible to combine some or all of the results if the cohorts were within the clinically-relevant ranges for this diagnostic.

From these data, one quickly identifies a relatively large fraction of **negative** control samples considered, and a small number of positive control samples. In **Table 2A**, without positive controls we could not evaluate the performance because there were no TP values. However, it would be appropriate to incorporate these data in the second and/or third cohort because this data represents a significant set of negative controls.

In **Table 2B**, there are TP and TN for all taxa. The majority of organisms have performance metrics approaching 1, generally indicating strong performance around these organisms; however, the assay may warrant further investigation because 7 taxa show precision values of 0.5. Additional positive controls may result in improved performance values; updating the database could change the rate of FP; or possibly this could indicate a set of organisms that pose a significant challenge to this analysis, requiring additional confirmation.

And in **Table 2C**, 4 species had no TP. This is reflected in the sensitivity and precision of these organisms and demonstrates the potential value this approach generates assisting rapid problem identification. If another round of analyses with similar spike-in amounts of DNA were performed that gave TP values, then a full evaluation could be performed.

While the authors were clearly focused on specificity with these particular experiments, it should also be clear that using unbalanced sample sets and examining one aspect of the performance metrics can cause problems for overall evaluation.

### Further Perspective

One objection to this approach, given the large number of organisms to test, might be that it is intractable to test every single organism in a database to realize the full potential of mNGS analyses. We do not aim to render large portions of databases unusable, but some testing is necessary to evaluate any taxon’s fitness for purpose within an application. To alleviate some of the burden, we propose that *in silico* data, if vetted properly (such as a minimum of 3 datasets from independent sources) and at levels mimicking clinical samples, should inform on the analysis performance and significantly reduce the experimental burden on developers while improving the breadth of organisms that can be interrogated. Perhaps datasets could be generated, updated, and validated that are suitable for this purpose. Potentially, positive test results coming from purely *in silico* validated organisms could be flagged for additional analysis such as further bioinformatics, serotyping, or PCR.

As with any diagnostic test, the rigor necessary to attain regulated use is a practical reality, and with this work we have sought a measurement process capable of meeting these stringent requirements. As analyses achieve regulated use status the field will continue to collect data and examine differences between *in silico* and physical sample validation and make a well-reasoned decision on their role in validating mNGS analyses.

## Conclusions

In response to the challenge put forth from Chiu and Miller,^1^ criteria for evaluation of mNGS analyses were proposed that will improve evaluation of the analytical (and possibly clinical) validity of an mNGS-based analysis. We proposed an organism-centric reporting format that provides a clear summary of performance within a predefined cohort, and highlights strengths and weaknesses in an mNGS analysis and study design. The use of the HM and Pi to summarize each performance metric may enable developers and regulators to evaluate mNGS analysis performance and identify potential problems between two mNGS analyses, but such use should be reserved for comparison between similar data and cohorts.

## Acknowledgements

We would like to acknowledge Jayan Rammohan and Kevin Kiesler for helpful review and feedback. We also acknowledge funding from the US FDA CDRH.

